# The epitranscriptomic m6A RNA modification modulates synaptic function in ageing and in a mouse model of synucleinopathy

**DOI:** 10.1101/2024.09.12.612649

**Authors:** Avika Chopra, Mary Xylaki, Fanzheng Yin, Ricardo Castro-Hernández, Madiha Merghani, Valentina Grande, Brit Mollenhauer, André Fischer, Tgiago F. Outeiro

## Abstract

N6-methyladenosine (m6A) is the most abundant and conserved transcriptional modification in eukaryotic RNA, regulating RNA fate. While the functions of m6A in the development of the mammalian brain have been extensively studied, its roles in synaptic plasticity, cognitive decline, motor function, or other brain circuits remain underexplored. To date, the role of this modification in Parkinson’s disease (PD) and other synucleinopathies has been largely unknown. Here, we investigated the m6A epitranscriptome in a mouse model of synucleinopathy. We performed m6A RNA immunoprecipitation sequencing (meRIP-seq) to obtain the m6A epitranscriptome of the midbrain in young (3 mo) and aged (15 mo) A30P-aSyn transgenic mice (aSyn Tg) and C57BL6 control wild type (Wt) mice. We observed hypermethylation of synaptic genes in 3 mo aSyn Tg mice compared to age-matched Wt mice. This methylation was reduced during ageing, with synaptic genes becoming increasingly hypomethylated. Using immunofluorescence imaging alongside biochemical analysis, we further investigated the expression of m6A regulatory enzymes — writer, N6-Adenosine-Methyltransferase Complex Catalytic Subunit (METTL3); reader, YTH N6-methyladenosine RNA-binding protein (YTHDF1); and eraser, fat mass and obesity-associated protein (FTO) — in the cortex, striatum, hippocampus, and cerebellum of Wt and aSyn Tg mice, as well as in primary cortical neuronal cultures. We observed that the levels of METTL3, YTHDF1 and FTO were similar between Wt and aSyn Tg mice. Interestingly, the writer protein METTL3 was found in both the nucleus and in the post-synaptic compartment in neuronal cultures. Our findings suggest that alterations in the regulation of m6A RNA methylation may be associated with neurodegeneration and ageing and that this level of epitranscriptomic regulation plays a significant role at the synapse.

## Introduction

Epigenetics is a process that acts as a bridge between genotype and phenotype, changing the outcome of a gene without modifying the DNA sequence. It encompasses various types of alterations such as methylation of DNA, histone modifications, and chromatin remodeling [1,2]. The field that studies posttranscriptional changes of RNA bases is known as epitranscriptomics and aims to identify modifications in the transcriptome and unravel their regulatory roles [3]. N^6^-methyladenine (m6A) modification is the most prevalent mRNA modification occurring at a frequency of 0.15 - 0.6% of all adenosines [4–6] and is found in different types of RNAs including protein coding, messenger RNA (mRNA) [7,8]. This modification is being heavily studied due to its prevalence and importance in eukaryotes, where it regulates various cellular functions [8–12]. Methylation marks are mostly added co-transcriptionally to specific consensus motif sequences and are mostly found close to the 3’ un-translated region (3’-UTR) at the stop codon [4,13].

The m6A modification machinery consists of three groups of enzymes, namely methyltransferases, demethylases and detectors of this modification, also termed writers, erasers, and readers, respectively. The modification can be added onto the RNA by methyltransferases including methyltransferase-like 3 (METTL3), methyltransferase-like 14 (METTL14), and adapter protein Wilms tumour 1-associated protein (WTAP) [5,14–19]. METTL3 was the first discovered methyltransferase and was described as a key component in the methylation of transcripts involved in physiological processes such as embryonic development and brain development [20–23]. The catalytically active part of METTL3 forms a heterodimer with METTL14, which provides the structure to install the m6A modification onto RNA [5,24–27]. Apart from these, the m6A methyltransferase complex comprises other proteins, including RNA binding proteins (RBP) like RBP15, which recruit this machinery to the target mRNA to facilitate methylation [15,28]. Demethylases, for instance, fat mass and obesity-associated protein (FTO) and alkB homologue 5 (ALKBH5) remove the m6A modification marks making this a dynamic process [29,30]. FTO was the first eraser protein identified [29,31] and it carries out oxidative demethylation where m6A is converted in several steps to N^6^-hydroxymethyl adenosine (hm6A), then to N^6^-formyladenosine (f6A) and lastly to adenine [31]. Additionally, reader proteins like the YT521-B homology (YTH) domain family of proteins including YTHDF1, YTHDF2/3, insulin-like growth factor 2 mRNA-binding proteins (IGF2BP1), heterogeneous nuclear ribonucleoprotein (HNRNPA2B1) recognise the modification and modulate alternative splicing and processing of target transcripts [8,32]. The ones located in the cytoplasm include YTHDF1, YTHDF2, and YTHDF3. The main function of these proteins is unclear; however, studies indicate different roles of these enzymes in a cell. While YTHDF1 promotes the translation of mRNA thereby promoting its protein expression, YTHDF2 promotes the degradation of mRNA, and YTHDF3 promotes both translation and degradation of mRNA [33,34].

m6A methylation has been associated with ageing, and downregulation in the expression of FTO and an overall increase in RNA m6A modification was found in human follicular fluid (FF) and granulosa cells (GCs) from aged donors and murine ovaries from old mice [35]. Another recent study has also highlighted the relationship between elevated m6A levels and brain ageing in both mice and humans [36], although a significant downregulation of m6A methylation in ageing mice and in postmortem brain tissue from Alzheimer’s disease patients has also been reported [37].

Strikingly, ageing is also a major risk factor for the onset and progression of neurodegenerative disorders such as Parkinson’s disease (PD) and related synucleinopathies, disorders associated with the accumulation of aggregated alpha-synuclein (aSyn) protein in the brain. Despite the growing interest in the role of m6A methylation in ageing, its role in PD is still unclear and remains controversial. While a recent study indicated that the expression levels of m6A are significantly reduced in cellular models of PD and in the striatum of a rodent model of PD [38], a different study showed that the m6A methyl transferases were significantly decreased while the demethylases were increased in the substantia nigra and in the striatum of mouse models of PD [39]. Ageing also affects multiple homeostatic pathways including mitochondrial function and protein degradation pathways, ultimately leading to cell death in the substantia nigra [40–42].

In this study, we investigated the m6A RNA modification profile in the context of ageing and neurodegeneration in a mouse model of synucleinopathy. We performed methylated RNA immunoprecipitation sequencing (meRIP-seq) to obtain the m6A epitranscriptome of the midbrain of C57BL6 (Wt) control and A30P aSyn PD (aSyn Tg) mice at 3 months (3 mo) and 15 months (15 mo) of age. We observed an increase in the m6A methylation in the genes involved in the molecular and biological processes at the synapse. Several genes involved in synaptic processes were hypermethylated in 3 mo aSyn Tg vs. Wt mice, while the methylation marks were reduced in 15 mo aSyn Tg vs. age-matched Wt mice. Using immunofluorescence imaging of primary cortical neurons, we observed that the localisation of the writer METTL3 enzyme is not only limited to the nucleus but is also present at the post-synapse. Our findings suggest that m6A RNA methylation is possibly playing a role in neurodegeneration and ageing, and is involved in regulating synaptic function.

## Results

### m6A alterations during ageing in Wt mice and in a transgenic model of synucleinopathy

We began by examining the distribution of differentially expressed transcripts and transcripts with m6A methylation in the healthy ageing brain of Wt mice. We further extended the analysis to aSyn Tg mice, in order to compare the effects of ageing on m6A modification and assess this change in the context of synucleinopathy. We extracted and dissected the midbrains from five 3 mo and five 15 mo Wt and an equal number of aSyn Tg mice, as this brain region is particularly relevant in the context of PD and related synucleinopathies. Methylated RNA immunoprecipitation sequencing (meRIP-seq) was performed to determine the midbrain-specific epitranscriptomic landscape in 3 mo and 15 mo mice (Figure 1A, B). First, we compared the differential expression of genes due to ageing in 15 mo vs 3 mo mice in both Wt and aSyn Tg animals. We observed a stronger effect in aSyn Tg mice where 270 transcripts were differentially expressed when compared with Wt mice, where only 79 transcripts were differentially expressed. Most differentially expressed transcripts in aSyn Tg mice were downregulated (Supplementary Figure S1C). However, when we compared the differentially methylated transcripts, we found similar numbers between Wt (420 genes) and Tg (424 genes) mice (Figure 1A).

**Figure 1.**
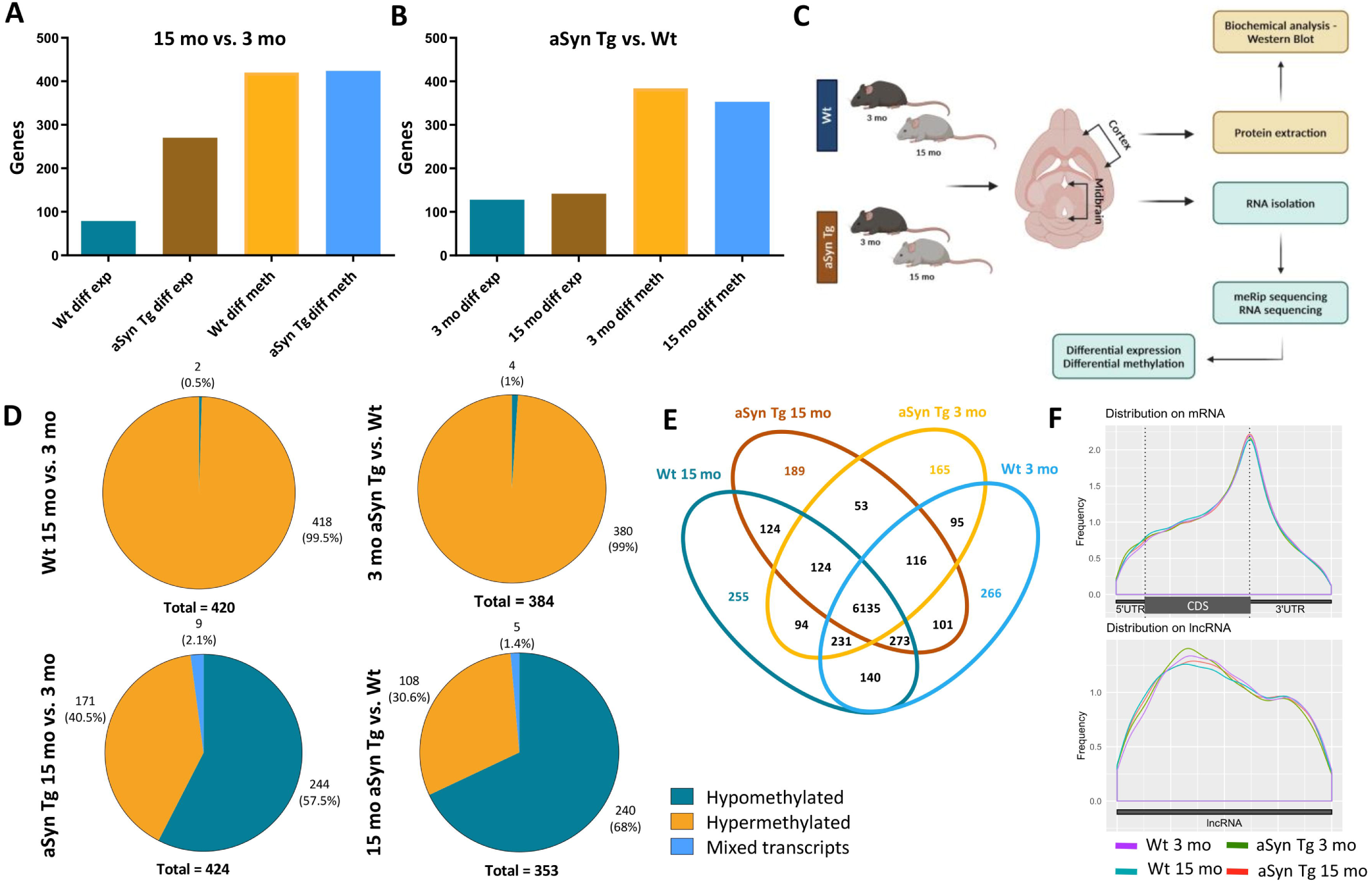
m6A modification landscape in ageing and aSyn Tg mice. (A) Bar plots representing the variations in the differentially expressed genes and differentially methylated genes in the midbrain from aSyn Tg mice than Wt mice when 3mo mice are compared with 15mo applying equal cutoffs for FC and adjusted P value (FC > 1.2 and padj ≤ 0.05). Left: with age, right: with aSyn (B) Bar plots representing the effects of aSyn pathology on differential expression and differential methylation in 3mo and 15mo Wt vs. aSyn Tg mice. (C) Overview of protocols utilized in this project. (D) Pie charts representing the overview of the distribution of hypomethylated, hypermethylated and mixed transcripts as an effect of aSyn pathology and ageing. (E) Venn diagram representing the number of differentially methylated genes involved in young and old Wt and aSyn Tg mice. It also provides a summary of the overlapping genes across different genotypes. (F) Guitar plots representing the distribution of m6A methylation on mRNA and lncRNA. CDS stands for coding sequence, and UTR stands for untranslated region.

On the other hand, the effect of aSyn A30P overexpression was assessed by a comparison between aSyn Tg and Wt mice at 3 mo and 15 mo. We did not identify major differences in the number of differentially expressed transcripts which were 128 at 3 mo and 142 at 15 mo. Similarly, the differentially methylated transcripts showed no significant changes between Wt and aSyn Tg at 3 mo (384 genes) and 15 mo (353 genes) (Figure 1B). While we did not see a significant difference in the differential methylation, we found that the prevalence of hyper- and hypomethylated transcripts was reversed (Figure 1B).

As a consequence of ageing, the methylation levels of the differentially methylated m6A transcripts in 15 mo vs. 3 mo Wt mice we found 418 hypermethylated and only 2 hypomethylated transcripts. On the contrary, 171 genes were hypermethylated and 244 transcripts were hypomethylated in 15 mo vs. 3 mo aSyn Tg mice (Figure 1D). A transcript can carry multiple m6A methylation marks and some of these marks likely increase while others decrease simultaneously. Such transcripts are referred to as mixed transcripts. We found only 9 such mixed transcripts (Figure 1D, left column) in aSyn Tg mice during ageing.

To identify the effect of aSyn expression on the levels of the m6A modification, we compared 3 mo aSyn Tg mice with age-matched Wt mice. We found 380 hypermethylated transcripts and only 4 transcripts were hypomethylated, and no mixed transcripts. On the other hand, when comparing 15 mo aSyn Tg vs. Wt mice, a reverse trend was observed, with 108 hypermethylated, 240 hypomethylated, and only 5 mixed transcripts (Figure 1D, right column).

We found 6135 methylated transcripts in common between all groups (Figure 1E). The 3 mo Wt mice displayed 266 unique methylated transcripts and the 15 mo Wt mice 255. These numbers were lower for the aSyn Tg group where the 3 mo mice showed 165 unique methylated transcripts and the 15 mo mice showed 189. At 15 months, both genotypes showed 124 commonly methylated transcripts, suggesting the occurrence of conserved processes in normal ageing and synucleinopathies. The aSyn Tg mice displayed 53 methylated transcripts correlated with aSyn expression and conserved with ageing.

The overall distribution of m6A methylation peaks on mRNA did not display any striking anomalies in any of the groups. As expected, the majority of the m6A modifications were localised close to the 3’ UTR on mRNA (Figure 1F), as previously reported for ageing-related genes [36].

### m6A levels are reduced in synaptic transcripts upon ageing

To investigate the cellular processes and the subcellular regions where the differentially methylated transcripts were identified in 15 mo aSyn Tg vs. Wt mice, we performed gene ontology (GO) term analysis. We found that synaptic function was the biological process most represented. Our analysis revealed that transcripts with m6A methylation marks have strong enrichment for biological processes such as receptor internalization, excitatory post-synaptic potential, axonal transport, and dendritic spine morphogenesis. The identified transcripts were also involved in cellular processes including neuron projection cytoplasm and synaptic vesicle membrane (Figure 2A). Previous data from our group has also shown that while the average dendritic spine number was not significantly different, a strong difference was observed in the abundance of different types of spines in aSyn Tg neurons when compared with Wt neurons [67].

**Figure 2.**
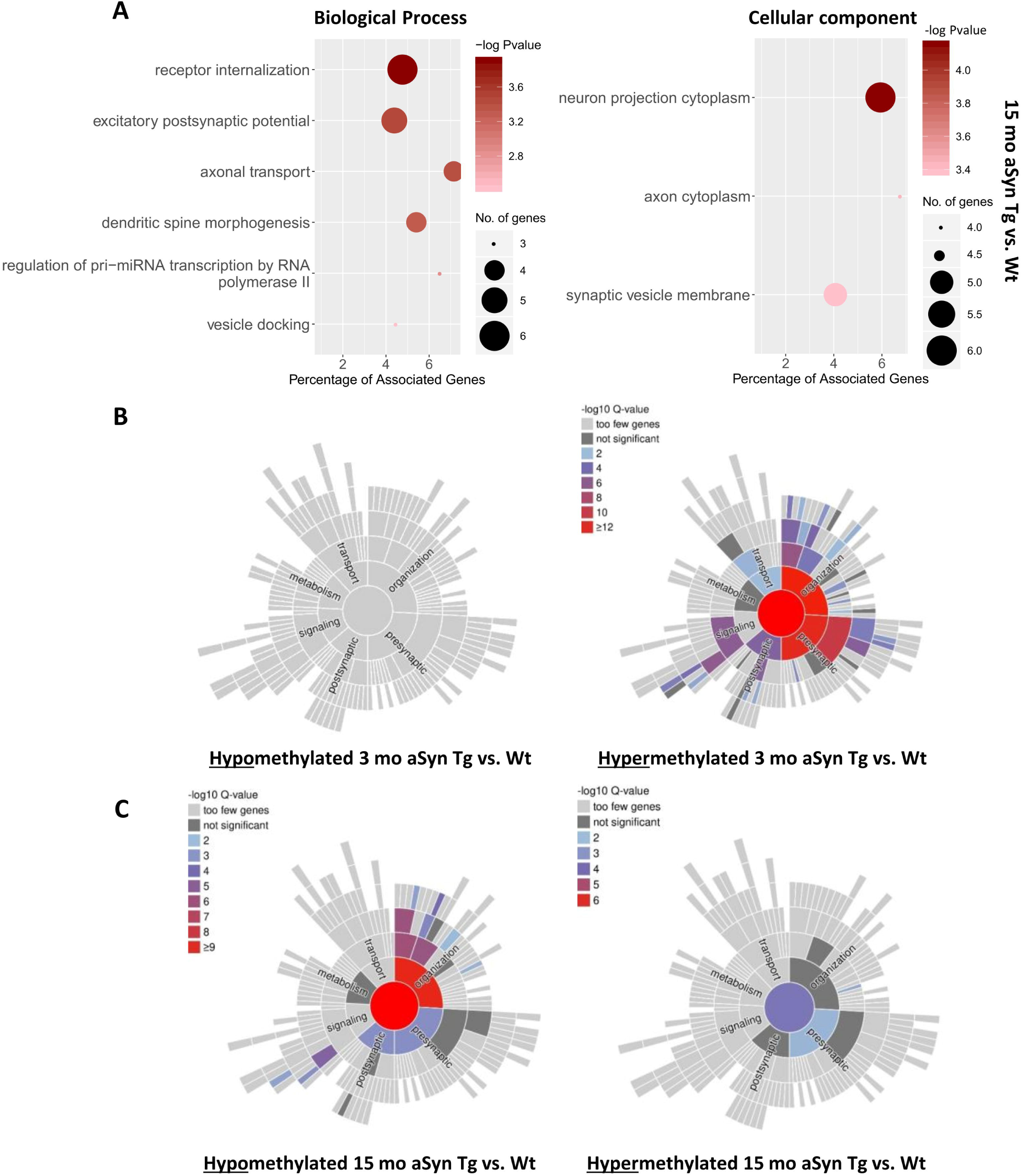
Gene ontology (GO) analysis for 3 mo and 15 mo Wt vs. aSyn Tg mice. (A) GO analysis for the biological processes and cellular processes in 15 mo aSyn Tg vs. Wt mice, FDR = 1.5. (B) Sunburst plots representing GO analysis for the genes involved in the processes at the synapse in aSyn Tg vs. Wt mice at 3 mo. Graphs created with syngoportal.org. (C) Sunburst plots representing GO analysis for the genes involved in the processes at the synapse in aSyn Tg vs. Wt mice aged mice at 15 mo. Graphs created with syngoportal.org.

We also observed that when we compared the differentially enriched genes in aSyn Tg 15 mo vs. 3 mo, we saw a strong enrichment of pathways involved in the immune system. The genes with highest enrichment score were involved in innate immune system, adaptive immune system, immunoglobulin binding, phagocytic vesicle, defense response, innate immune response, and regulation of immune system processes among others (Supplementary Figure S2).

We used the SynGO software [64] (https://syngoportal.org/), an experimentally annotated database for synaptic location and functional GO, to specifically check the m6A methylation of synaptic transcripts. These differentially methylated genes are primarily involved in synaptic organisation, metabolism, transport, signaling, and both pre-and post-synaptic processes. Based on meRIP and RNA sequencing analysis, sunburst plots of Wt and aSyn Tg mice were generated (Figure 2B). Our findings suggest that the differentially methylated transcripts that are involved at the synapse and in synaptic function were majorly hypermethylated in 3 mo aSyn Tg mice compared to 3 mo Wt mice (Figure 2A). In 3 mo mice, the hypermethylated transcripts were concentrated in synapse organisation and at the pre-synapse (Figure 2B). Conversely, at 15 months of age, the majority of the differentially methylated transcripts were hypomethylated when aSyn Tg mice are compared with Wt mice, which were largely involved in signalling, pre- and post-synapse, and synapse organisation (Figure 2C).

In summary, we observed that the differentially methylated genes, both hyper- and hypomethylated genes in 15 mo aSyn Tg vs. Wt mice were mostly hypomethylated and involved in synaptic processes, especially at the pre-and post-synapse. However, at 3 mo, most of the differentially methylated genes were hypermethylated.

### The levels of m6A regulatory proteins do not change in different brain regions in Wt and aSyn Tg mice

To validate the meRIP-Seq results and to evaluate if the overall levels of the m6A modification are affected in aSyn Tg vs. Wt mice, we performed an m6A ELISA, for total RNA isolated from cortical and midbrain brain tissue of young and aged, Wt and aSyn Tg mice. We found that overall, the levels of m6A modification marks were higher in the cortex compared to the midbrain. In cortical brain tissue, we observed that the m6A marks in the aSyn Tg mice do not display the same pattern as in Wt animals (Figure 3A). A significant reduction in m6A marks was observed in 12 mo aSyn Tg mice compared to 12 mo Wt mice (Wt; mean = 0.032 ± 0.018 OD, aSyn Tg; mean = 0.011 ± 0.002 OD, adjusted p-value = 0.045). In the younger animals, however not significant, a slight reduction in m6A marks was observed in 3 mo aSyn Tg were compared with 3 mo Wt mice, indicative of an effect that probably starts at a young age and progresses with ageing (Figure 3A). In the midbrain, we did not observe significant differences in the m6A marks between the groups (Figure 3A). This is consistent with the meRIP-Seq data where we observed that the total number of differentially methylated transcripts was not altered among the groups (Figure 1A, B). This suggests that despite the observed hypo- and hyper-methylation in certain transcripts (Figure 1D) the overall m6A levels remain in an equilibrium in the midbrain.

**Figure 3.**
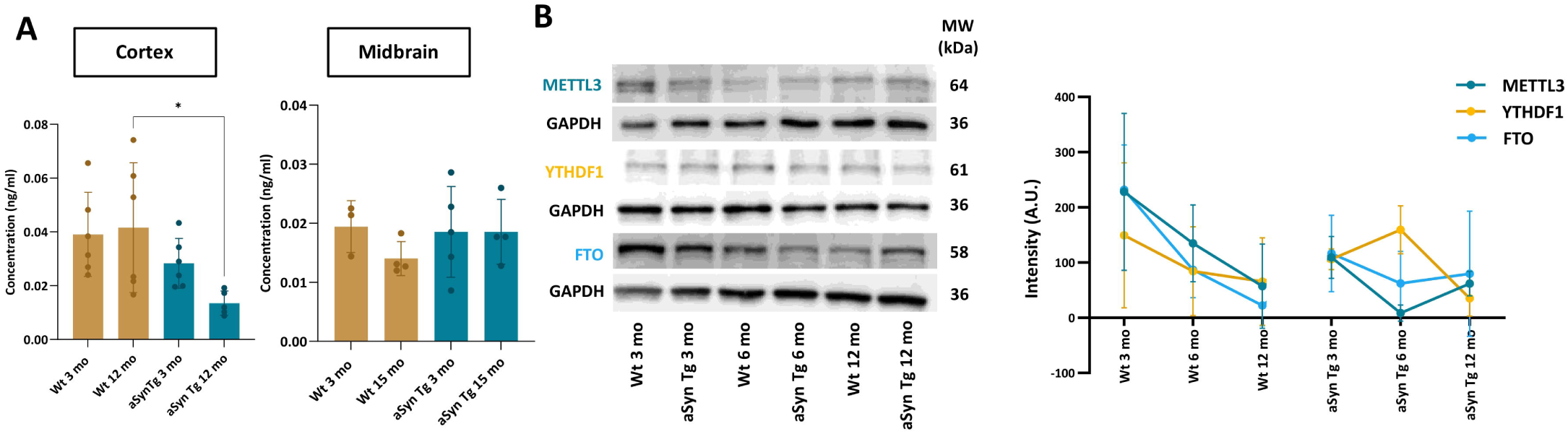
Levels of m6A RNA and m6A regulators in mouse brain tissue. (A) Bar plots representing the levels of m6A modification on total RNA using ELISA from cortical and midbrain regions of Wt and aSyn Tg mice. A significant reduction in the levels of m6A was observed in the aged, 12 mo aSyn Tg mouse cortex when compared with 12 mo Wt mice (*, ajd P value = 0.045). No significant differences were observed in the midbrain region of the brain. (B) Biochemical analysis using western blot comparing the expression levels of METTL3, YTHDF1 and FTO in the cortical tissue from 3 mo, 6mo and 12 mo Wt vs. aSyn Tg mice. Scatter plot showing no significant difference in the expression level when comparing Wt with aSyn Tg mice.

Next, we asked if the observed changes in the m6A marks in the cortex correlate with altered levels of the three m6A regulatory proteins in the cortical brain tissue. To do so, we performed immunoblotting analysis to assess the expression of m6A regulators — METTL3, YTHDF1, and FTO. We used three different age groups – 3 mo, 6mo and 12 mo from the Wt and aSyn Tg mice for our analysis. We observed no significant differences in the expression levels of these m6A regulatory proteins neither within genotypes (Tg vs. Wt) nor across the different age groups (Figure 3B). Nonetheless, we observed a trend towards a reduction in the levels of METTL3 and FTO in Wt mice during ageing (Figure 3B).

Considering no difference in the protein levels of m6A regulators, we sought to identify if the sub-regional and/or subcellular localisation of the three m6A regulatory proteins is altered. For this, we assessed the levels of METTL3, YTHDF1 and FTO by immunofluorescence intensity analysis in four different regions of the mouse brain, namely cortex, striatum, hippocampus, and cerebellum. MAP2 co-staining was used only to identify the neurons present in the areas of the brain. All these proteins were present in our selected areas of interest, but their compartmentalization varied across different regions of the neurons.

The m6A writer protein, METTL3 was observed as small puncta within the nucleus in the cortex, striatum, hippocampus, and cerebellum in both aged Wt and aSyn Tg brain (Figure 4A). We found no significant differences in protein levels between aSyn Tg vs. Wt mice based on the fluorescence intensity (Figure 4B).

**Figure 4.**
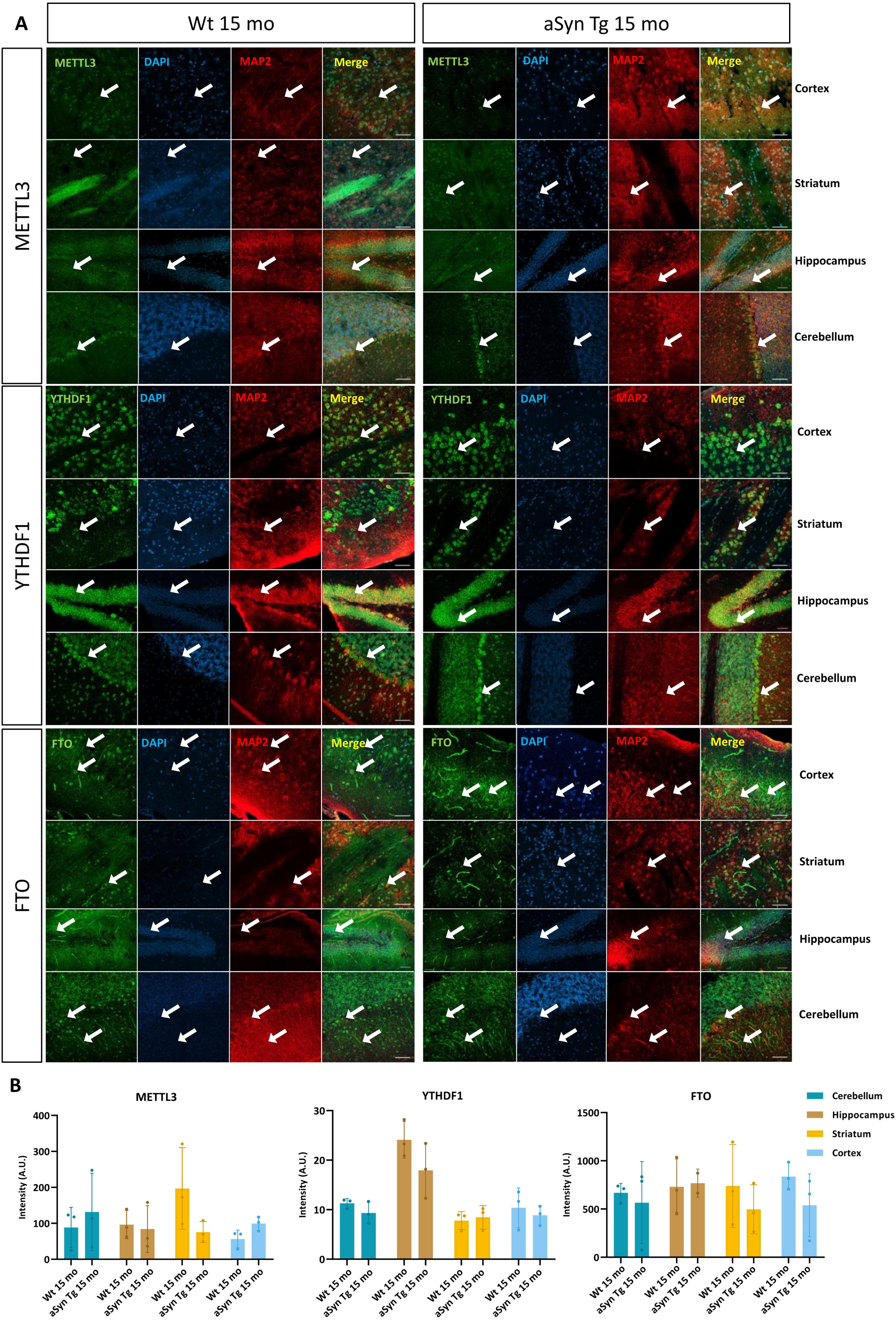
Distribution of m6A regulatory proteins METTL3, YTHDF1 and FTO in the cortex, striatum, hippocampus and cerebellum of mouse brain. (A) Representative images of mouse brain tissue slices (30µm) fixed, and stained with MAP2, METTL3, YTHDF1 and FTO. The images are acquired using a confocal microscope, 40x magnification with tiles, and scale bar = 50µm. White arrows point to the neurons with the protein expression. (B) Bar plots showing the comparison between the mean intensity of METTL3, YTHDF1 and FTO in 15mo Wt vs. aSyn Tg mice.

The m6A reader protein, YTHDF1, on the other hand, localised only in the soma of neurons in all four areas of interest, with a similar expression pattern in Wt and aSyn Tg mouse brain slices. (Figure 4A). No significant difference in the fluorescence intensity of the protein was observed in 15 mo aSyn Tg mice when they were compared with Wt mice (Figure 4B).

Finally, the m6A eraser, FTO, was detected in the nucleus and the soma of cells in the cortex, striatum, hippocampus, and cerebellum. FTO was also localized in the dendrites of neurons in the cortex and cerebellum in both Wt and aSyn Tg mouse brain slices (Figure 4A). After analysis of the expression of this protein in the soma, no significant differences were observed in the different regions of either aSyn Tg or Wt mice (Figure 4B).

Our findings show that the m6A regulatory proteins – METTL3, YTHDF1 and FTO are present in the cortex, striatum, hippocampus, and the cerebellum in the mouse brain. In addition, we found that the levels of METTL3, YTHDF1 and FTO do not change in cortical brain tissue.

### METTL3 is reduced in the aSyn Tg mouse post-synapse

Based on our bioinformatics analysis, we observed that the majority of m6a genes were synaptic, so we asked if/and to what degree the m6A regulators are present at synapses acting as local regulators since the overall regulator levels remain unchanged. We used primary cortical neurons that allow higher imaging potential and performed co-staining of the three m6A regulatory proteins – METTL3, YTHDF1 and FTO alongside Syn1, pre-synaptic marker and PSD95, post-synaptic marker to study the localisation of these proteins at the synapse. Previous studies have indicated reduced synaptic activity in neurons derived from sporadic PD patients when compared with controls [65,66].

Two experimenters performed the colocalisation analysis, and similar results were obtained. METTL3 was observed to be present in the nucleus as nuclear speckles and it significantly co-localized with PSD95, indicating its presence at the post-synapse in both Wt (PCC = 0.810 ± 0.039) and aSyn Tg (PCC = 0.748 ± 0.075) primary neurons. In contrast, METTL3 did not co-localize with synaptotagmin1 in Wt (PCC: 0.489 ± 0.086) or aSyn Tg (PCC: 0.476 ± 0.050) primary neurons, indicating no noticeable localisation at the pre-synapse (Figure 5B).

**Figure 5.**
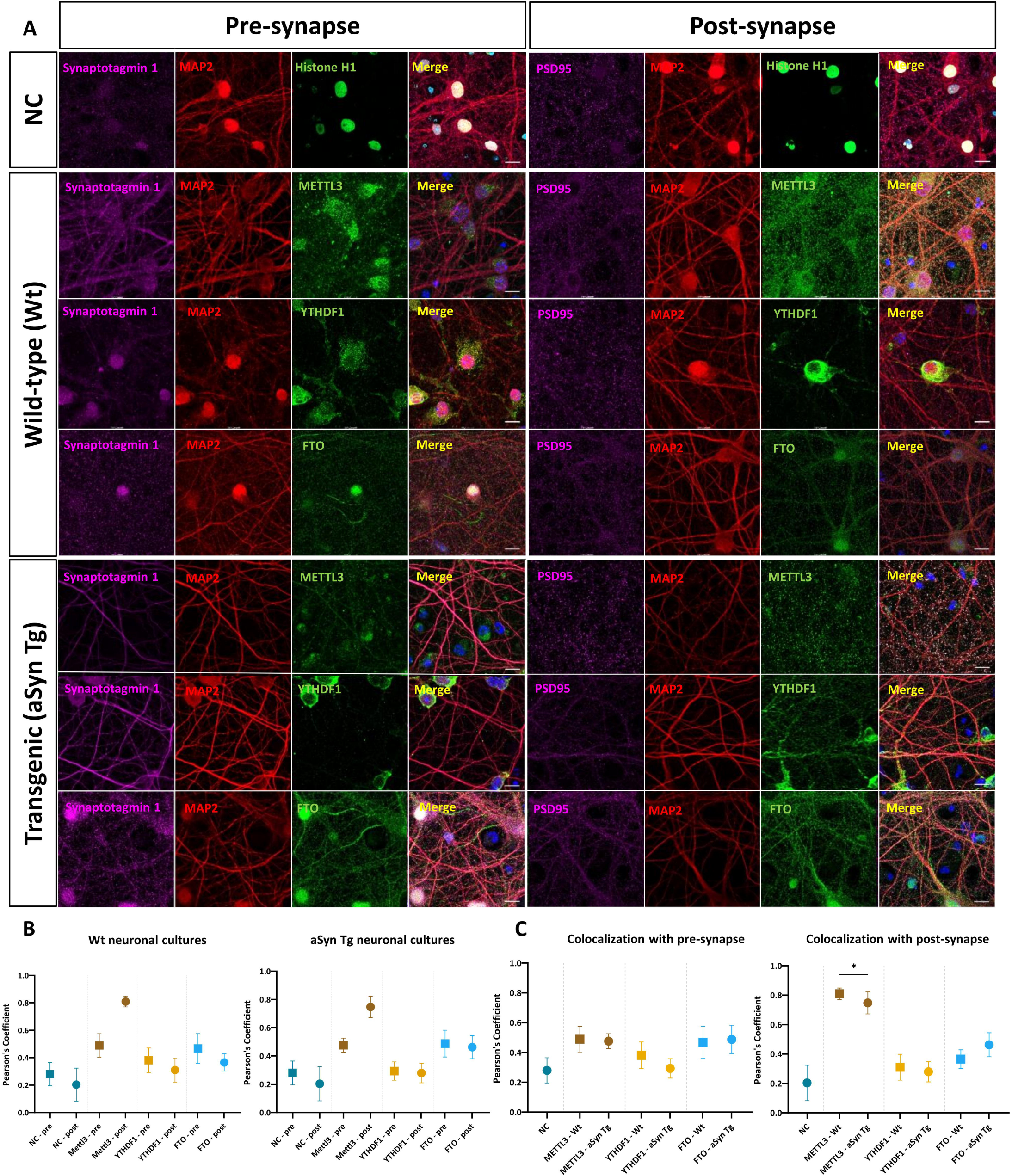
METTL3 is localised at the post-synapse in Wt and aSyn Tg DIV14 primary neurons. (A) Representative images of DIV14 primary neurons, fixed and stained with Histone H1, MAP2, PSD95, Synaptotagmin 1, METTL3, YTHDF1, and FTO in Wt and aSyn Tg neurons. Images are acquired using airy-scan at 63x magnification, scale bar = 10µm. (B) Box and whisker plot representing the colocalisation of m6A regulators in Wt and aSyn Tg D-14 primary neurons at the pre-and post-synapse. METTL3 colocalises with the post-synapse, (Pearson’s correlation coefficient, Wt = 0.810, aSyn Tg = 0.748). A significant difference (*, adj p value = 0.018) was observed between METTL3 at the post-synapse in Wt neurons when compared with aSyn Tg neurons. (C) Box and whisker plot representing the colocalisation of m6A regulators at the pre-and post-synapse in Wt and aSyn Tg primary neurons. METTL3 colocalisation is significantly reduced in aSyn Tg neurons when compared with Wt at the post-synapse

Similarly, we investigated the compartmentalization of YTHDF1, which did not overlap with either synaptotagmin1 (PCC: Wt = 0.382 ± 0.090, aSyn Tg = 0.294 ± 0.065) or PSD95 (PCC: Wt = 0.310 ± 0.088, aSyn Tg = 0.280 ± 0.069) (Figure 5B). The m6A eraser protein, FTO, was identified within the cell nucleus and in some cases, in the soma of primary neurons. No significant co-localisation of FTO was observed at the pre-(PCC, Wt = 0.468 ± 0.109, aSyn Tg = 0.488 ± 0.095) or post-synapse (PCC, Wt = 0.366 ± 0.063, aSyn Tg = 0.463 ± 0.081) (Figure 5B).

Overall, we observed that the expression of METTL3 was reduced at the post-synapse in aSyn Tg primary neurons suggesting a possible loss of METTL3 at the synapse.

### Primary neurons from aSyn Tg mice have a greater number of synapses than Wt animals

Since we observed a reduction in the localisation of METTL3 at the synapse in aSyn Tg primary neurons, we wondered whether this has any influence on the synapse morphology and number, or vice versa. For this, we used primary cortical neurons and performed proximity analysis of the pre-and post-synaptic markers Syn1 and PSD95 using the IMARIS software Imaris (RRID: SCR_007370) and the data were analysed blindly to avoid bias. We found that the number of synapses in aSyn Tg primary neurons was significantly increased when compared with Wt neurons (Wt; mean = 2607 ± 763.1, aSyn Tg; mean = 3152 ± 1187, p-value = 0.044) (Figure 6).

**Figure 6.**
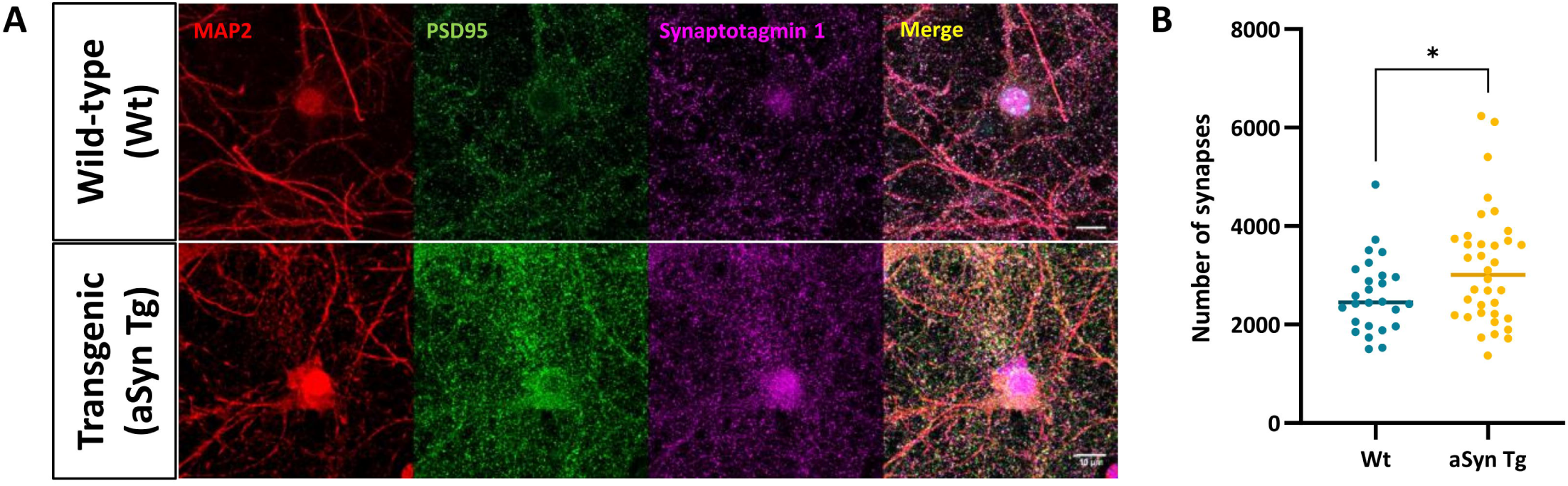
The number of synapses is increased in aSyn Tg primary neurons. . (A) Representative immunofluorescent images of DIV14 primary neurons stained with pre-(Syn1, magenta) and post-synaptic (PSD95, green) markers and MAP2 in red. Acquired at 63x magnification, airy scan mode. Scale bar = 10µm. (B) Scatter plot for synapse counting for Wt and aSyn Tg mouse models with a significant reduction in the number of synapses (*, adj p value = 0.044) in aSyn Tg primary neurons when compared with Wt.

## Discussion

The involvement of epitranscriptomics in PD is still not well understood. In this study, our objective was to begin to unravel the role of m6A RNA methylation in the A30P aSyn Tg mouse model of PD/synucleinopathy, in disease and physiological ageing.

Our findings revealed that 99.5% of the transcripts were hypermethylated in 15 mo vs. 3 mo Wt mice suggesting that ageing might contribute to a significant increase in m6A methylation. This observation aligns with a recent study that has demonstrated an increase in m6A methylation during ageing in both mice and humans [36]. This indicates that the accumulation of m6A methylation marks might be a general feature of ageing in the brain, potentially influencing various biological processes. It has also been shown that these epigenetic changes arise mainly in young and aged mice, while during the adolescent stage these genes are stabilised [68]. In contrast, when comparing 15 mo aSyn Tg mice with their 3 mo counterpart littermates, more transcripts were hypomethylated. This decrease in methylation with age in aSyn Tg mice might suggest that the loss of methylation might be related to aSyn expression/pathology. There is a possibility that the overexpressed aSyn is somehow interacting with the m6A regulators and inhibiting methylation. Previous studies have shown that the 6-hydroxydopamine rat PD model has significantly reduced m6A levels in the striatal brain region and in the cellular model of PD [69]. Similarly, a reduction in the total levels of m6A was also reported in the MPTP-treated PD mouse model and in MPP+-treated MN9D cells [70].

The effect of aSyn expression was more directly studied in the comparison between 3 mo aSyn Tg and Wt mice. The increase in methylation suggests that aSyn expression may be linked to enhanced methylation activity in early life. However, in 15 mo aSyn Tg vs. Wt mice, we observed more hypomethylated genes, suggesting that A30P aSyn expression, and likely pathology, might interfere with the writing or erasing of methylation marks during ageing. Importantly, our results are in accordance with a recent publication suggesting that the levels of m6A are significantly lower in PD patients [71].

Interestingly, based on GO term analysis, we found that differentially methylated transcripts were predominantly associated with synaptic function and that methylation was lost with ageing. Hypermethylation of transcripts involved in the synaptic processes might lead to an inhibition of mRNA translation and thereafter reduced functionality of the synapse. This is particularly relevant as more than half of the causative genes and risk factors in PD have been identified to function at the synapse [72] and this change in methylation pattern might indirectly affect the functioning of the synapse. Considering that different RNA-based epigenetic mechanisms occur locally at the synapse, this underscores their potential importance in synaptic regulation [74]. Additionally, GO term analysis of the differentially expressed transcripts showed a strong enrichment in immune response-related pathways. Notably, m6A is implicated in the regulation of inflammation under physiological and pathological conditions, known as “inflammatory ageing”, or inflammaging. This is a chronic inflammatory state arising during ageing and it is characterised by an increase in pro-inflammatory factors such as tumor necrosis factor (TNF-α), interleukins (IL-1β and IL-6), and C-reactive protein (CRP), to name just a few [73].

The assessment of total m6A methylation marks showed a significant reduction in the cortex of 12 mo aSyn Tg mice when compared with age-matched Wt mice. This suggests that the overall m6A methylation decreases in the ageing brain. This finding is consistent with recent studies that reported a predominant reduction in m6A methylation in the frontal cortex and cingulate cortex from PD patients [71,75]. In vitro studies have also reported a significant reduction in m6A levels in the cellular PD models [70]. Despite this, we observed similar expression levels of m6A regulatory proteins. However, our findings are consistent with those from a recent study reporting that the levels of YTHDF1 and FTO mRNA remain unchanged in PD. However, a reduction in the levels of METTL3 mRNA in PD patients was also reported in that study, in contrast to what we observed [71]. Some studies have also reported up-regulation of ALKBH5, IGF2BP2 in the substantia nigra while, YTHDF1 and fragile X mRNA (FMR1) were downregulated. In the striatum, an up-regulation of FMR1 and Cbl proto-oncogene like 1 (CBLL1) and down-regulation of METTL3, RBM15 and IGF2BP3 was observed [76].

Although we found no significant differences in the levels of m6A regulatory proteins, we observed that the three proteins were present in the cortex, striatum, hippocampus, and cerebellum. Our findings are consistent with those from previous studies reporting that the m6A-methyltransferase complex, including METTL3, is predominantly found in the nucleus, but that it can also be localised in the cytoplasm [77–79]. As methylation generally occurs co-transcriptionally in the nucleus, while reading occurs largely at the cytosol, YTHDF1 is primarily cytosolic [80]. FTO, on the other hand, is localised both in the nucleus and in the cytosol [81]. These observations suggest that an inhibitory factor affecting their function and aSyn aggregation is potentially playing a role. Previous studies have shown that aSyn binds to histone proteins, which may alter the protein surface due to post-translation modifications of histone tails, altering their affinity for DNA and other histones [82]. Another study showed that aSyn expression increases histone-H3 lysine-9 (H3K9) methylation, suggesting a possible interaction of aSyn with epigenetic writer proteins [83]. This could constitute a mechanism through which aSyn affects gene methylation, ultimately affecting synaptic functioning [84,85].

Given that most of the hypermethylated genes were synaptic, we assessed the presence of these regulators at the synapse. Our findings suggest the presence of METTL3 at the post-synapse, indicating that the enzyme might be locally active at the synapse. The dynamic nature of m6A methylation may be localised to different regions within a cell, depending on the temporal requirements of the respective genes [86,87]. In another study, m6A regulatory proteins were found localised adjacent to the synapse and in the dendrites, also suggesting the sub-cellular localisation of m6A methylated mRNA which might lead to synaptic modifications [88,89]. Our findings revealed that the expression of METTL3 is reduced at the post-synapse in aSyn Tg DIV14 neurons, suggesting a possible loss of METTL3 at the post-synapse. Due to this reduction at an early stage, the nucleus might be sending hypermethylated transcripts to the dendrites to compensate for this loss. However, this may be lost during ageing. Based on our bioinformatics analysis, we observed that methylated transcripts associated with synaptic function were reduced in aSyn Tg mice with ageing. This suggests that a reduction in the m6A methylation in synaptic genes may be directly linked with the reduced METTL3 localisation at the synapse in aSyn Tg neurons.

## Conclusion

In conclusion, our study highlights the critical role of m6A RNA methylation in the progression of PD and its implications in ageing, particularly focusing on the A30P aSyn mouse model. Our findings indicate that aSyn expression/pathology may influence RNA methylation patterns, leading to reduced levels of m6A in mice. The differentially methylated transcripts were predominantly synaptic, indicating that m6A modifications play a crucial role in synapse physiology and potentially contribute to the synaptic dysfunction observed in PD [90]. Although we observed no significant changes in the levels of m6A regulatory proteins, the reduction in METTL3 at the synapse in aSyn Tg mice informs on the importance of subcellular compartmentalization of these proteins in the regulation of RNA methylation. Therefore, our research provides new insights into the epigenetic mechanisms underlying PD and underscores the need for further exploration of m6A RNA methylation as a potential therapeutic target for PD and other related synucleinopathies.

## Materials and methods

### Resource availability

#### Lead contact for reagent and resource sharing

Further information and requests for resources and reagents should be directed to and will be fulfilled by the lead contact, Tiago F. Outeiro.

### Experimental model and subject details

#### Animals

Five male homozygous [A30P] aSyn transgenic (aSyn Tg) mice (C57BL/6J-Tg(Th-SNCA*A30P*A53T) 39Eric/J, Strain #:008239, RRID: IMSR_JAX:008239) and five wild type (Wt) littermates, 3 mo, 6mo, 12 mo and 15 mo were included in this study for RNA sequencing and analysis. Additional animals used for each experiment are mentioned in the respective section. Animals were housed in standard cages with 12h light/12h dark cycle and ad libitum access to food and water in the animal facility of the University Medical Centre Göttingen (Göttingen, Germany). Animal procedures were performed in accordance with the European Community (Directive 2010/63/EU); and institutional and national guidelines under the Lower Saxony State Office for Consumer Protection and Food Safety (LAVES) (license number 19.3213). Animals were anesthetised with carbon dioxide (CO_2_) and then sacrificed using cervical dislocation. For tissue processing, the brain was surgically extracted from the skull and was further dissected to isolate the cortex and midbrain. The tissue was snap-frozen in liquid nitrogen and preserved at -80°C until use.

For the preparation of brain slices, the animals were perfused with 1x phosphate buffer saline (PBS) followed by 4% paraformaldehyde (PFA). The whole brain was dissected out of the skull and placed overnight in 4% PFA at 4°C. The following day, the brain was washed once with 1x PBS and then placed in 15% saccharose and 30% saccharose with 0.1% sodium azide (NaN_3_). The brains were placed at 4°C until used. Sagittal cryo-sections of 30 µm thickness were cryo-sectioned using Leica CM3050 S cryostat for each mouse brain. The sections were placed in 1x PBS with NaN_3_ until they were used for immunohistochemistry (IHC) protocol.

#### Neuronal cultures

Primary cortical neuronal cultures were prepared from both aSyn Tg and Wt mice as previously described (Saal et al., 2015). Cells were seeded on coverslips coated with poly-L-ornithine (0.1 mg/mL in borate buffer; PLO) (Sigma-Aldrich, MO, USA) or culture plates (Corning, Merck, Darmstadt, Germany) for immunocytochemistry and maintained in neuronal cell culture medium (Neurobasal medium, Gibco Invitrogen, CA, USA), supplemented with 1% penicillin-streptomycin (PAN Biotech, Aidenbach, Germany), 0.25% GlutaMAX (Gibco Invitrogen, CA, USA), and 2% B27 (B-27™ Plus Supplement (50X), Gibco, #A3582801). The cells were maintained at 37°C with 5% CO_2_, and one-third of the medium was replaced with fresh media every 3-4 days.

#### Immunocytochemistry

Primary neurons were first washed with 1x PBS (PAN Biotech, Aidenbach, Germany) and fixed with 4% of PFA for 20 min at room temperature (RT). Cells were first permeabilized with 0.1% Triton X-100 (Sigma-Aldrich, MO, USA) for 10 min and then blocked with 1.5% normal goat serum (NGS) in 1x PBS (Sigma-Aldrich, MO, USA) for 1h at RT. The cells were then incubated with the primary antibodies diluted in blocking solution (FTO (mouse, Abcam – ab92821, 1:1000), METTL3 (rabbit, Cell Signalling - #51104, 1:1000), YTHDF1 (rabbit, Proteintech 17479-1-AP, 1:1000), MAP2 (rabbit, Proteintech – #17490-1-AP, 1:1000), postsynaptic density protein 95 (PSD95) (mouse, Synaptic systems 124011, 1:1000), PSD95(D74D3) (rabbit, Cell Signalling - #3409, 1:1000), synaptotagmin 1 (Syn1) (mouse, Synaptic systems - #105311, 1:1000) synaptotagmin1 (rabbit, Synaptic systems – 105102/3, 1:1000)) overnight at 4°C, with constant shaking at a speed of 40-50 rpm. The following day, cells were washed three times for 10 min each with 1x PBS (PAN Biotech, Aidenbach, Germany) and then incubated with fluorescently conjugated secondary antibodies (Alexa Fluor 488 donkey anti-mouse IgG – Invitrogen, Alexa Fluor 488 donkey anti-rabbit IgG - Invitrogen, Alexa Fluor 555 donkey anti-mouse IgG – Invitrogen, Alexa Fluor 555 donkey anti-rabbit IgG – Invitrogen, Alexa fluor 633 goat anti-chicken IgG - Invitrogen) for 2h at RT, on a shaker at 40-50 rpm. Lastly, nuclei were counter-stained with DAPI (Carl Roth, Karlsruhe, Germany) and mounted on glass slides (Epredia™ SuperFrostPlus™ Adhesion) with mounting media (Invitrogen™ Fluoromount-G™) for microscopy.

#### Immunohistochemistry

The free-floating tissue was carefully transferred from PBS and NaNH_3_ to 1x tris-buffered saline (TBS) in a new 24-well plate with immunohistochemistry (IHC) grids. The tissue was washed three times for 5 min each with 1x TBS to remove the remnants of NaN_3_. Following this, tissue antigen retrieval was performed using citrate buffer (10 mM citrate, 0.05% Tween 20, pH 6.0) at 90°C for 30 min in a water bath. The tissue samples were allowed to cool down until they reached RT. Tissue sections were washed thrice for 10 min each with 1x TBS and then incubated with 0.3% Triton X-100 for 20 min. Without washing, the tissue slices were incubated directly in a blocking solution (5% BSA, and 1% NGS in 1x TBS) for 2h at RT. The tissue sections were then incubated for 48h with antibodies diluted 1:200 times (FTO (mouse, Abcam – ab92821, 1:1000), METTL3 (rabbit, Cell Signalling - #51104, 1:1000), YTHDF1 (rabbit, Proteintech #17479-1-AP, 1:1000), MAP2 (rabbit, Proteintech – #17490-1-AP, 1:1000)) at 4°C, on a shaker. The following day, tissue sections were washed thrice for 10 min each with 1x TBS at RT. Afterwards, the tissue sections were incubated in secondary antibody (1:400 dilution) (Alexa Fluor 488 donkey anti-mouse IgG – Invitrogen, Alexa Fluor 488 donkey anti-rabbit IgG - Invitrogen, Alexa Fluor 555 donkey anti-mouse IgG – Invitrogen, Alexa Fluor 555 donkey anti-rabbit IgG – Invitrogen, Alexa fluor 633 goat anti-chicken IgG - Invitrogen) overnight at 4°C on the shaker. The following day, the tissue was washed thrice for 10 min each with 1x TBS to remove the unbound secondary antibody. The tissue sections were incubated for 5 min in DAPI (Carl Roth, Karlsruhe, Germany) followed by a 10 min wash with 1x TBS. The tissue sections were then mounted onto glass slides (Epredia™ SuperFrostPlus™ Adhesion) and left to dry in the dark at RT. 8-10µl of mounting media (Invitrogen™ Fluoromount - G™) was applied as a drop onto each section on the glass slide and a glass coverslip (Avantor, 631-0147, 0.16-0.19mm thickness, #1.5) was carefully placed on the tissue to avoid any air bubbles.

#### Western blots

Cortical brain tissue was extracted from three adult mice per age group (age group 2 mo, 6 mo and 12 mo) from each Wt and Tg mice. The tissue from adult mice was lysed using homogenisation beads (1.4 mm diameter ceramic beads, 91-PCS-CK14B, PEQLAB Biotechnologie, Germany) in RIPA buffer (50 mM Tris, pH 8.0, 0.15M NaCl, 0.1% SDS, 1.0% NP-40, 0.5% Na-Deoxycholate, 2 mM EDTA) supplemented with protease and phosphatase inhibitors cocktail (cOmplete™, Mini Protease Inhibitor Cocktail, Roche). The homogenisation was performed in Precellys 24 tissue homogenizer (Bertin Instruments) at 65 000 × g for 2 × 30 sec runs with a 30 sec break in between. Protein concentrations in the lysates were determined by the BCA protein assay (Pierce Biotechnology, #23225) following the manufacturer’s protocol. The spectrophotometric recordings were performed using a plate reader (Tecan, Infinite® M200 pro) at 562 nm. Volume for 20 μg of protein per sample was calculated, to which Laemmli sample buffer (250 mM Tris-HCl pH 6.8, 10% SDS, 1.25% Bromophenol Blue, 5% ß-mercaptoethanol, 50% glycerol) was added (1:5 dilution of Laemmli: protein). The proteins were denatured for 5 min at 95°C, with 600 rpm in a thermal shaker. The samples were loaded on a 12% SDS-PAGE electrophoresis gel and separated at 120V for approximately 1h. The proteins were then transferred to PVDF membranes (iBlot2 Transfer Stack, PVDF, Life, #IB24002X3) using iBlot2 (Invitrogen, CA, USA) using the P0 protocol for 7 min. Membranes were blocked with 5% BSA (Sigma Aldrich, #AG9418) in TBS (pH 8) with 0.05% Tween-20 (TBS-T) and then incubated with the appropriate primary antibody (FTO (mouse, Abcam – ab92821, 1:5000), METTL3 (rabbit, Cell Signalling - #51104, 1:1000), YTHDF1 (rabbit, Proteintech #17479-1-AP, 1:2500)) overnight in 5% BSA (Sigma-Aldrich, MO, USA) in TBS-T at 4°C. After three washes with TBS-T, membranes were incubated for 2h with horseradish peroxidase (HRP) conjugated secondary antibodies (Cytiva sheep anti-mouse IgG, 1:8000 dilution, Cytiva donkey anti-rabbit IgG, 1:8000 dilution). Following incubation, membranes were washed three times with TBS-T for 10 min each and developed in a chemiluminescence system (Fusion FX Vilber Lourmat, Vilber, France) using chemiluminescent HRP substrate (Millipore, MA, USA). Intensities of specific bands were normalised to a protein loading control.

#### RNA extraction

For brain tissue, TRIzol reagent (Invitrogen, CA, USA) and ceramic beads (1.4 mm diameter ceramic beads, 91-PCS-CK14B, PEQLAB Biotechnologie, Germany) were added in vials with frozen tissue and homogenisation was performed with the Precellys 24 tissue homogenizer (Bertin Instruments) at 65 000 g for 2 x 30 sec runs with a 30 sec break in between. RNA isolation was performed with TRIzol reagent according to the manufacturer’s instructions. Briefly, 0.5 ml of TRIzol was added to the tissue samples and incubated for 5 min before homogenisation. After homogenisation, 0.1 ml of chloroform was added to each tube. The tubes were shaken vigorously and incubated at RT for 2-3 min. The tubes were centrifuged at 14 000g, for 10min at 4°C. The top aqueous layer was carefully transferred to a fresh tube. Total RNA was then precipitated using Clean and Concentrator 5, Zymo research kit (Zymo Research, #R1014). Following RNA extraction and precipitation, RNA was dissolved in RNAse-free water and concentration and quality were estimated with NanoDrop 1000 spectrophotometer (Thermo Fisher Scientific, MA, USA) by measuring absorption at 260 nm and the ratios 260/280 nm and 230/280nm respectively.

#### m6A fluorescence ELISA

m6A RNA methylation ELISA (m6A RNA methylation, Abcam, #ab233491) was used for the detection of m6A RNA methylation in the cortical brain and midbrain tissue from 3 mo and 12 mo/15 mo Wt and aSyn Tg mice. For this experiment, we used cortical mouse brain tissue from Wt mice each at 3 mo and 12 mo (six 3 mo and six 12 mo), and aSyn Tg mice (six 3 mo and five 12 mo). Midbrain tissue was used from Wt mice (three 3 mo and four 15 mo) and aSyn Tg mice (five 3 mo and four 15 mo). This protocol ensures that the total m6A methylation is measured in both mRNAs and small RNAs. The protocol was performed according to the manufacturer’s protocol. Briefly, total RNA isolated from the tissue was added to bind to the assay wells. The wells were then washed to remove the unbound RNA and diluted capture antibody was added to the wells. Unbound capture antibody was washed out with multiple washing steps. Diluted detector antibody was then added along with fluoro-enhancer. The wells were washed again and incubated with fluoro developer mixture to initiate the fluorescence reaction. The fluorescence was then measured at 590nm using plate reader (Tecan, Infinite® M200 pro).

#### MeRiP

Total RNA was treated with DNase to remove any DNA remnants and further, 5-10 µg of total RNA was depleted for rRNA using the RiboMinus Eukaryote Ribosomal Removal Kit (Invitrogen). The large RNA fraction (>200 bp) was fragmented to a fragment size of approximately 100-120 nt by RNA Fragmentation Reagent at 70°C for 5 min (#AM8740, Thermo Fisher Scientific, Waltham, MA). 5% of this RNA sample was kept as inputs and the rest was subjected to immunoprecipitation (IP). 300 µg of fragmented RNA was used for each IP. 3 µg of anti-m6A antibody was incubated with the RNA IP buffer (0.2 M Tris-HCl pH 7.5, 0.5 M NaCl and 2% (vol/vol) Igepal CA-630) for 2h at 4°C with constant rotation. RNA antibody mixes were then crosslinked twice with 0.15 J/cm2 of UV light (254 nm) in a UVP crosslinker (Analitik Jena). Antibody-RNA conjugates were incubated with 30 μl of Protein A/G beads overnight (ON) at 4°C with rotation in 500 μl IP buffer supplemented with 200 units. The beads with immunoprecipitated RNA were washed with IP buffer five times, and further washed with low-salt (50 mM Tris pH 7.4, 50 mM NaCl, 1mM EDTA, 1, 0.1% NP-40, 0.1% SDS) and high-salt (same as low-salt but with 500mM NaCl) buffers at 4°C with rotation to remove any nonspecific binding. RNA was eluted by incubating in 150 μl PKD buffer (Qiagen) with 10 μl of Proteinase K (Millipore) for 1 hour at 37°C with agitation. Eluted RNA was cleaned before proceeding to library preparation.

#### Library Preparation and Sequencing

Samples were prepared for sequencing using the SMARTer Stranded Total RNA-Seq Kit v2 — Pico Input Mammalian (Takara) according to the manufacturer’s protocol. All the RNA obtained from the IP samples was used for library preparation, for input samples, 2 ng was used. The libraries were amplified for a total of 12 cycles for input and 16 cycles for IP samples. RNA-Seq samples were prepared with the TruSeq Stranded mRNA Library Prep Kit (Illumina) according to the manufacturer’s instructions for which, 50 ng of rRNA-depleted RNA from each of the inputs was used. Prepared libraries were sequenced in a Hiseq 2000 System (Illumina) for 50 cycles in single-end reads. Additional metadata is also available via the GEO database (GSE275432).

#### Bioinformatic Analysis of meRIP-seq and RNA-seq

Raw reads were processed and demultiplexed using bcl2fastq (v2.20.2), and low-quality reads were filtered out with Cutadapt v1.11.0 [43]. Filtered reads were mapped to the mouse (mm10) genome using the STAR aligner v2.5.2b [44]. The resulting bam files were sorted and indexed, and the unmapped reads were removed using SAMtools v1.9.0 [45]. Methylation sites were determined using MeTPeak v1.0.0 [46], and differential methylation (hypo- and hyper-methylated regions) was assessed with ExomePeak v2.16.0 [47] using 3 mo samples as control and 15 mo as treatment for the first comparison and for the second comparison, Wt samples were used as controls while aSyn Tg were used as treatment. An adjusted P value (padj, also termed as FDR [false discovery rate]) cutoff of 0.05 and FC cutoff of 1.2 or 1.5 were used as indicated in the text. Only consistently significantly, differentially methylated peaks were used, unless indicated.

For RNA-seq analyses, read counts were obtained with subread’s featureCounts v1.5.1 [48] from the bam files of input samples. Differential gene expression was determined by DESeq2 v3.5.12 [49] using normalised read counts and correcting for covariates detected by RUVseq v1.16.1 [50]. Cutoffs of padj ≤ 0.05, FC ≥ 1.2, and BaseMean ≥ 50 were applied to the results (Supplementary Figure S1A, B). Background expressed genes were determined for each region as those genes with a BaseMean > 50 in the corresponding input sample.

For visualization, bam files of both IP and input samples were collapsed for PCR duplicates using SAMtools, and IP samples were normalized to their corresponding inputs and to their library size using deeptools v3.2.1 [51] bamCompare. The resulting normalized tracks were visualized in the IGV Browser 2.9.2 [52].

#### GO Analysis

GO term enrichment analyses were performed using the App ClueGO v2.5.3 [53] in Cytoscape 3.7.2 [53], with GO Term Fusion enabled to collapse terms containing very similar gene lists and using a custom background corresponding to expressed genes in the corresponding species as obtained from RNA-seq results of the corresponding input samples of the meRIP experiments. GO term tables for biological processes, cellular components, pathways, and KEGG were produced and are labelled accordingly in the figures. The resulting enriched GO terms were visualized with a custom script using ggplot2 v3.3.5 [54] playing the adjusted p-value (padj) for the GO term, the number of genes from the list that belong to said term, and the percentage of the total genes in the GO term that are present in the list. Synaptic GO enrichment analyses were performed with SynGO (v1.1, syngoportal.org) [55].

#### Additional Bioinformatics Packages and Tools

Scripts and analysis pipelines were written in R (3.5.2) [56]. Peak annotation was performed with Homer v4.10.4 [57] and Annotatr v1.8.0 [58]. Guitar plots were produced with the Guitar v1.20.1 [59] R package. Volcano plots were generated with plot.ly/orca v4.9.4.1 [60] (Supplementary Figure S1C). Area-proportional Venn diagrams were produced with BioVenn (www.biovenn.nl), and multiple list comparisons performed with Intervene / UpSet (asntech.shinyapps.io/intervene/). Mouse/human homologues were determined by their annotation in NCBI’s HomoloGene database using the HomoloGene (v1.4.68.19.3.27) R package. Odds ratios and p values to determine significance in overlapped datasets were calculated with the GeneOverlap R package v1.18.0 [61]. De novo motif analyses were performed with Homer’s findMotifsGenome, and motifs containing the DRACH consensus sequence out of the top 10 most significant are dipcasplayed. KEGG pathway enrichment was produced with KEGG Mapper (www.genome.jp/kegg/mapper/) [62]. Microscopy images were pre-processed with Fiji, and quantification was automated in Cell Profiler (cellprofiler.org) [63]. Graphs, heat maps, and statistical analyses were performed on GraphPad Prism version 9.3.1 for Mac.

#### Microscopy

Images of the immunofluorescence-stained neurons (three experiments in total, with at least 10 regions of interest acquired per experiment) and brain tissue were acquired using a confocal microscope ZEISS LSM900. The images for the neuronal cell culture were acquired as a z-stack using 63x magnification with Airyscan mode. Brain slices were imaged at 40x magnification, with tiles in confocal mode.

#### Statistical analysis

The cell culture images for m6A regulators and their colocalisation at the synapse were analysed using FIJI with the plugin JaCoP, while the synapse counting was performed using IMARIS. IMARIS analysis was performed following the protocol from the Queensland Brain Institute. The protein signals were further classified inside the dendritic or outside using membranes based on the MAP2 signals. The signals were then divided into four groups, representing PSD95 inside the neuron, PSD95 outside the neuron, Synaptotagmin1 (Syn1) inside the neuron and Syn1 outside the neuron. When the signal for PSD95 inside the neuron was at < 500µm from the signal for Syn1, it was considered a synapse. Colocalisation of synaptic markers with m6A regulators was estimated using Pearson’s correlation coefficient (PCC) estimated using the JaCoP plugin in ImageJ, FIJI. All IHC images were also analysed for their expression levels using FIJI software. Data analysis was performed using the GraphPad software version 9 for Windows (GraphPad Software, La Jolla California USA, https://www.graphpad.com). For group comparisons, one-way ANOVA with Tukey’s test was used, while comparisons of two groups of means were done with an unpaired student’s t-test. All data are expressed as mean ± SD. Differences are considered significant with p1<10.05 (*p1<10.05, **<0.01, ***<0.001).

## Declarations

### Ethics approval

Animal procedures were performed in accordance with the European Community (Directive 2010/63/EU); and institutional and national guidelines under the Lower Saxony State Office for Consumer Protection and Food Safety (LAVES) (license number 19.3213).

### Consent for publication

All authors approved the publication of the article.

### Availability of data and materials

The datasets used and/or analysed during the current study are available from the corresponding author upon reasonable request. All raw data files for meRIPseq are available on the GEO database (GSE275432).

### Authors’ contributions

AC performed, analysed, and interpreted the data from cell cultures, ELISA, IHC, and western blot, and interpreted the meRIP results. MM and VG performed experiments in animals. MX interpreted the data for meRIP analysis and was a major contribution in writing the manuscript. YF performed and analysed ICC data. RCH performed the meRIP and subsequent data analysis. AC and TO wrote the manuscript. All authors read and approved the final manuscript.

## Supplementary Figures

**Figure S1. Gene expression analysis in aSyn Tg and Wt mice.** (A) Principle component analysis (PCA) plot of total Wt vs. aSyn Tg mice (PC1 = 40.5%, PC2 = 19.2%). (B) PCA plot of aSyn Tg – 3 mo and 15 mo mice, showing a clear segregation (PC1 = 49.8%, PC2 = 23.6%). (C) Volcano plot showing that the expression of most of the genes is down regulated in aSyn Tg mice, due to ageing (FDR < 0.05, abs(logFC) >1), aSyn Tg 15 mo vs. 3mo.

**Figure S2. Differentially expressed gene pathways in aSyn Tg 15 mo vs. 3 mo.** GO term analysis comparing the enrichment of pathways, molecular function, cellular components and biological processes of differentially methylated genes. Most pathway enrichment was observed in immune system and immune response linked pathways including – immunoglobulin binding, phagocytic vesicle, defense response and innate and adaptive immune response, FC > 1.2, FDR = 1.5.

