## Supplementary figures and images for "The epitranscriptomic m6A RNA modification modulates synaptic function in ageing and in a mouse model of synucleinopathy"

### Supplementary Figure 1

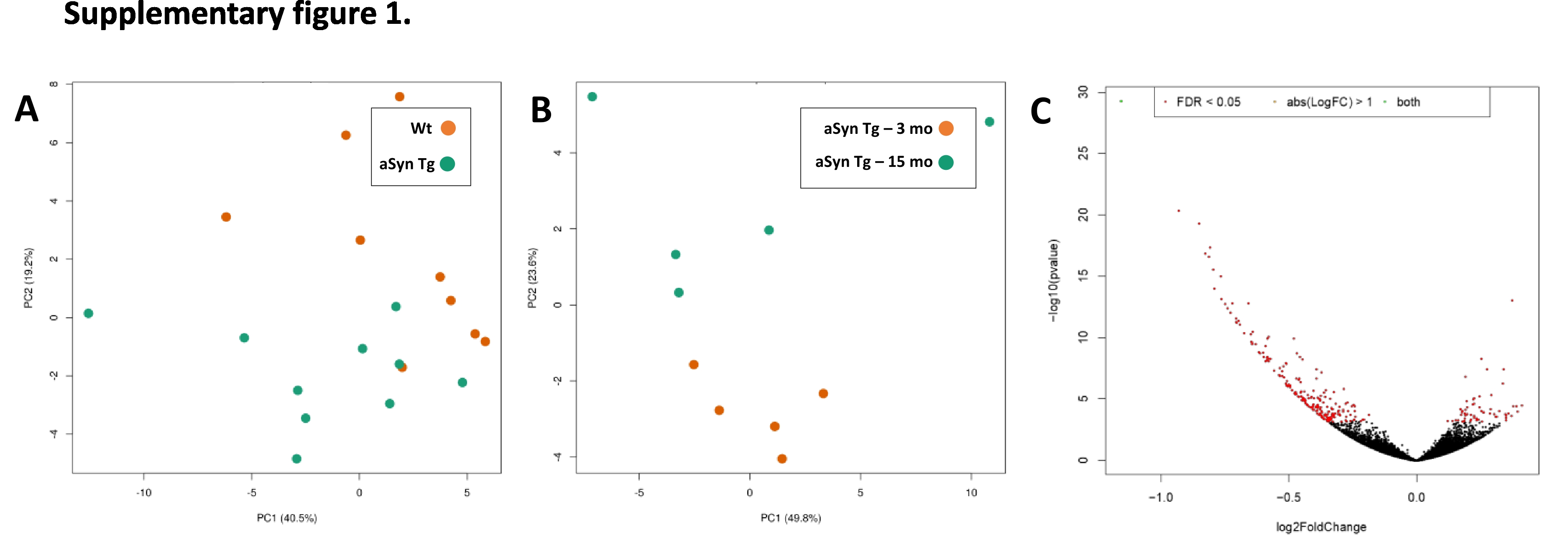

### Supplementary Figure 2

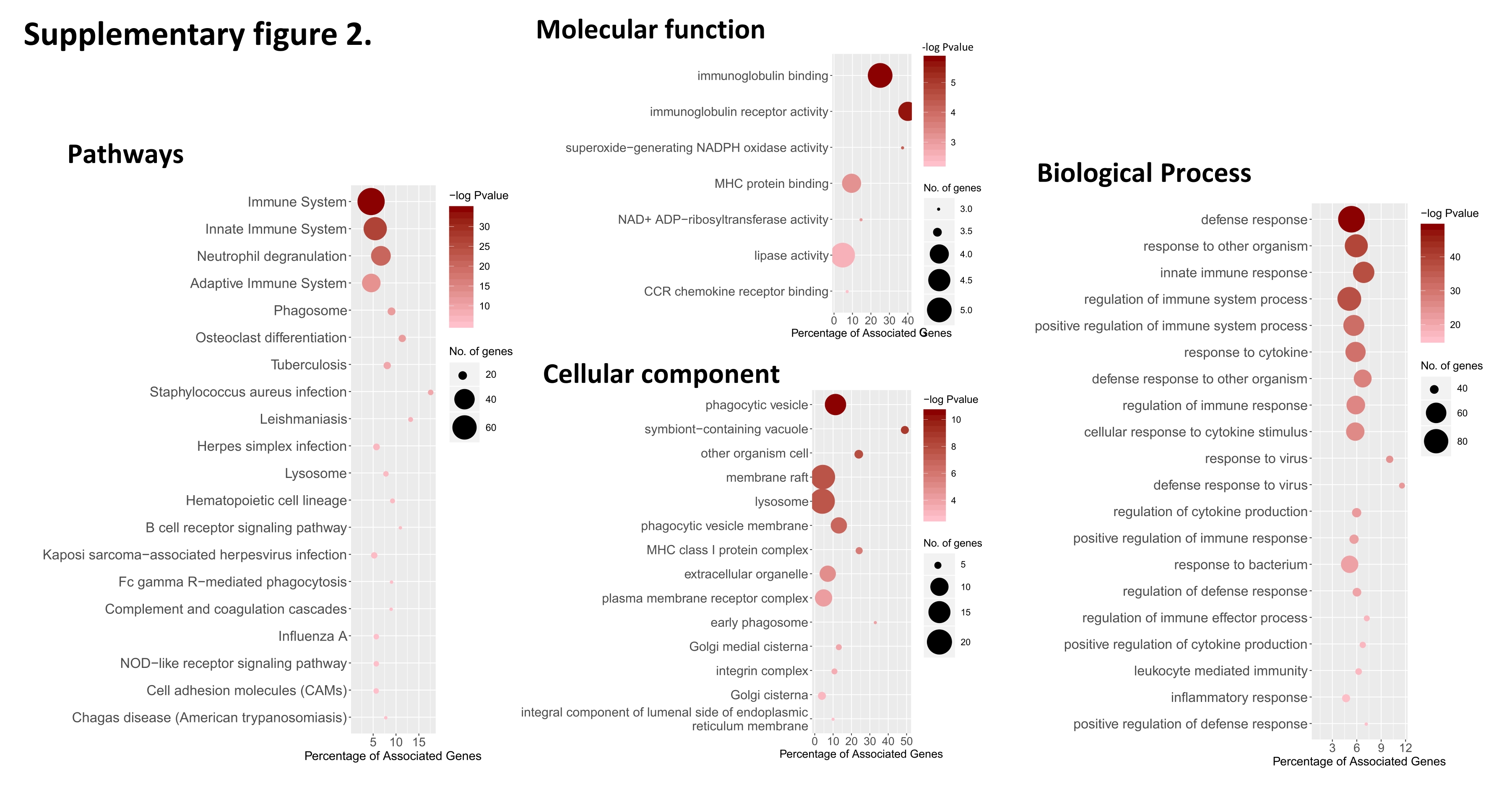
